# Modeling tumor dynamics and predicting response to chemo-, targeted-, and immune-therapies in a murine model of pancreatic cancer

**DOI:** 10.1101/2025.01.03.631015

**Authors:** Krithik Vishwanath, Hoon Choi, Mamta Gupta, Rong Zhou, Anna G. Sorace, Thomas E. Yankeelov, Ernesto A.B.F. Lima

## Abstract

We seek to establish a parsimonious mathematical framework for understanding the interaction and dynamics of the response of pancreatic cancer to the NGC triple chemotherapy regimen (mNab-paclitaxel, gemcitabine, and cisplatin), stromal-targeting drugs (calcipotriol and losartan), and an immune checkpoint inhibitor (anti-PD-L1). We developed a set of ordinary differential equations describing changes in tumor size (growth and regression) under the influence of five cocktails of treatments. Model calibration relies on three tumor volume measurements obtained over a 14-day period in a genetically engineered pancreatic cancer model (KrasLSLG12D-Trp53LSLR172H-Pdx1-Cre). Our model reproduces tumor growth in the control and treatment scenarios with an average concordance correlation coefficient (CCC) of 0.99±0.01. We conduct leave-one-out predictions (average CCC=0.74±0.06), mouse-specific predictions (average CCC=0.75±0.02), and hybrid, group-informed, mouse-specific predictions (average CCC=0.85±0.04). The developed mathematical model demonstrates high accuracy in fitting the experimental tumor data and a robust ability to predict tumor response to treatment. This approach has important implications for optimizing combination NGC treatment strategies.

## 1 Introduction

Pancreatic cancer, infamous for its aggressive growth, early metastasis, and resistance to conventional therapies, necessitates innovative treatment approaches [1, 2, 3]. Emerging treatment protocols are increasingly focused on strategies that target not just the cancer cells, but also the tumor microenvironment and immune system [4, 5, 6]. Pancreatic tumors, known for their dense stromal tissue and immunosuppressive environments, present challenges for traditional chemotherapies like cisplatin and gemcitabine [7, 8, 9, 10, 11, 12]. To address this, stromal-targeting agents such as calcipotriol and losartan are being investigated to enhance drug delivery, while immunotherapies, such as anti-PD-L1 (anti-Programmed Cell Death Ligand 1), are being tested to activate the immune system against the tumor [5, 13, 14, 15]. However, determining the optimal combination of these therapies and predicting tumor response remains an active area of research.

The literature on mathematical modeling in pancreatic cancer is sparse, especially when compared to other cancers [16, 17]. However, there are seminal efforts to model the efficacy of chemotherapy [18, 19] and immunotherapy [20, 21] in pancreatic cancer. In [18], Lee et al. examined the inconsistent responses of pancreatic cancer to gemcitabine observed between the in vitro (high efficacy) and in vivo settings (low efficacy). They used a system of partial differential equations to model cell proliferation, apoptosis, and nutrient diffusion gradients influenced by the microenvironment (e.g., inefficient vascularization or abundant stroma). The model, calibrated with in vitro data on drug exposure and cell viability and validated through in vivo mouse experiments, indicates that pancreatic tumors essentially resist gemcitabine due to poor vascularization. The study shows that the drug’s efficacy in controlled in vitro conditions, which mimic a single cell layer near the vasculature with optimal access to oxygen and nutrients, does not capture the complex dynamics of in vivo environments. In another work, Jenner et al. [19] used a hybrid agent-based model to evaluate the ability of locally delivered gemcitabine to treat pancreatic cancer effectively. The model was calibrated using both in vitro (drug release kinetics and cytotoxicity against human pancreatic cancer cells from gemcitabine-loaded alginate fibers) and in vivo (including tumor growth rates and responses from mouse experiments) data. In their approach, the authors specifically investigated how the tumor microenvironment, including factors like drug diffusion and cell proliferation rates, impacts the effectiveness of drug delivery. Their findings demonstrated that intratumoral placement of drug-loaded alginate fibers, which accounts for these microenvironmental factors, significantly improved treatment efficacy compared to peritumoral placement. They found that the drug release rate and pattern—such as constant, exponential, and sigmoidal releases—significantly influenced the drug’s ability to maintain therapeutic concentrations within the tumor microenvironment over time, with the exponential profile proving more effective in reducing tumor growth than others. This indicates that fine-tuning the release profile could be critical for optimizing treatment responses in pancreatic cancer.

Building on the investigation of drug delivery strategies within the complex pancreatic tumor microenvironment, other researchers have focused on understanding the interactions between pancreatic cancer cells and the immune system, which are crucial for improving patient survival. A system of five ordinary differential equations (ODEs) was developed by Hu et al. to investigate the dynamics of pancreatic cancer cells, their interactions with the immune system, and how this impacts patient survival [21]. This system models the interactions between pancreatic cancer cells, stellate cells, effector cells, and both tumor-promoting and tumor-suppressing cytokines. The model, which integrates 23 parameters sourced from the literature, was validated using survival data from two clinical trials. Findings based on optimal control theory indicated that mono-immunotherapy alone cannot effectively control pancreatic cancer, suggesting the necessity for combined therapies, including anti-TGF-*β* treatments and adoptive transfers of immune cells (a process where immune cells are harvested, sometimes genetically modified, and then infused back into the patient to boost the immune response), to enhance patient survival. Bratus et al. also employed an ODE framework to simulate the dynamics of cancer cell mutations and their interactions with CD8 T cells and nutrients [20]. The model describes the temporal dynamics of various pancreatic cancer cell populations differentiated by specific genetic mutations and includes mutation and fitness landscape matrices that characterize the cells’ survival capabilities. The model was able to effectively predict the growth dynamics of pancreatic cells with different mutations and their response to immune cells, notably demonstrating the effectiveness of immune cells in reducing tumor size. The authors did note, however, that incorporating experimental data into their study would improve the model’s validity.

In this contribution, our goal is to develop a mathematical framework capable of simulating and predicting the response of pancreatic tumors to various combination treatment regimens. While previous studies have provided valuable insights into the interactions between pancreatic cancer cells, the immune system, and drug delivery within the tumor microenvironment, they often focus on specific aspects, such as modeling the mutation dynamics within pancreatic cancer cells. Building on these foundational efforts, our work seeks to provide a comprehensive model that integrates the dynamics of chemotherapy, stromal-targeting drugs, and immunotherapy. To our knowledge, this represents the first effort to mathematically model the dynamics of NGC chemotherapy effects on pancreatic cancer. Specifically, we aim to model the dynamics of tumor growth and regression in response to combinations of chemotherapy, stromal-targeting drugs, and immunotherapy. By integrating experimental data from longitudinal tumor volume measurements obtained in murine models of pancreatic cancer, we calibrate our mathematical model to accurately capture the complex interactions between tumor cells, stromal components, and immune cells. We further demonstrate the ability of this model to predict patientspecific response to treatments. This study contributes to a biology-based, mathematical model that simulates complex treatment interactions and predicts responders and non-responders, which is crucial for optimizing personalized treatment strategies in pancreatic cancer.

## 2 Methods

### 2.1 Experimental Design

All animal procedures were approved by the institutional animal care and use committee (IACUC) of the University of Pennsylvania. The experimental procedures for acquiring the tumor data are detailed more thoroughly in [22].

#### 2.1.1 Genetically Engineering Model (GEM)

The mouse model employed for these studies was a genetically engineered model (GEM) of pancreatic ductal carcinoma (KrasLSLG12D-Trp53LSLR172H-Pdx1-Cre; usually referred to as KPC mice [23, 24, 25]) bred at the Mouse Hospital of Abramson Cancer Center of the University of Pennsylvania. Both the male and female KPC mice were enrolled. Once their tumor reached 50-150 mm^3^ estimated by MRI, mice were randomized and assigned to one of the treatment groups described below.

#### 2.1.2 Treatment

The treatment employed a chemotherapy backbone collectively referred to as NGC, consisting of Nab-paclitaxel (33 mg/kg), gemcitabine (266 mg/kg), and cisplatin (8 mg/kg for males and 4 mg/kg for females), stroma-directed drugs including calcipotriol (60 µg/kg) and losartan (30 mg/kg) and anti-PD-L1 mAb (200 µg/mouse). For the Nab-paclitaxel, murine albumin was used to develop murine albumin paclitaxel nanoparticles [26]. Six treatment groups were studied: control (untreated, *n* = 8), NGC (*n* = 18), NGC + losartan (*n* = 10), NGC + calcipotriol (*n* = 9), calcipotriol (*n* = 4), and NGC+ calcipotriol + immunotherapy (anti-PD-L1 mAb; *n* = 8). All drugs were administered via intraperitoneal injection under isoflurane anesthesia, and all mice were euthanized by cervical dislocation under anesthesia on day 14 following MR imaging as described in [22]. The dosing schedule for each drug is described in Figure 1.

**Figure 1.**
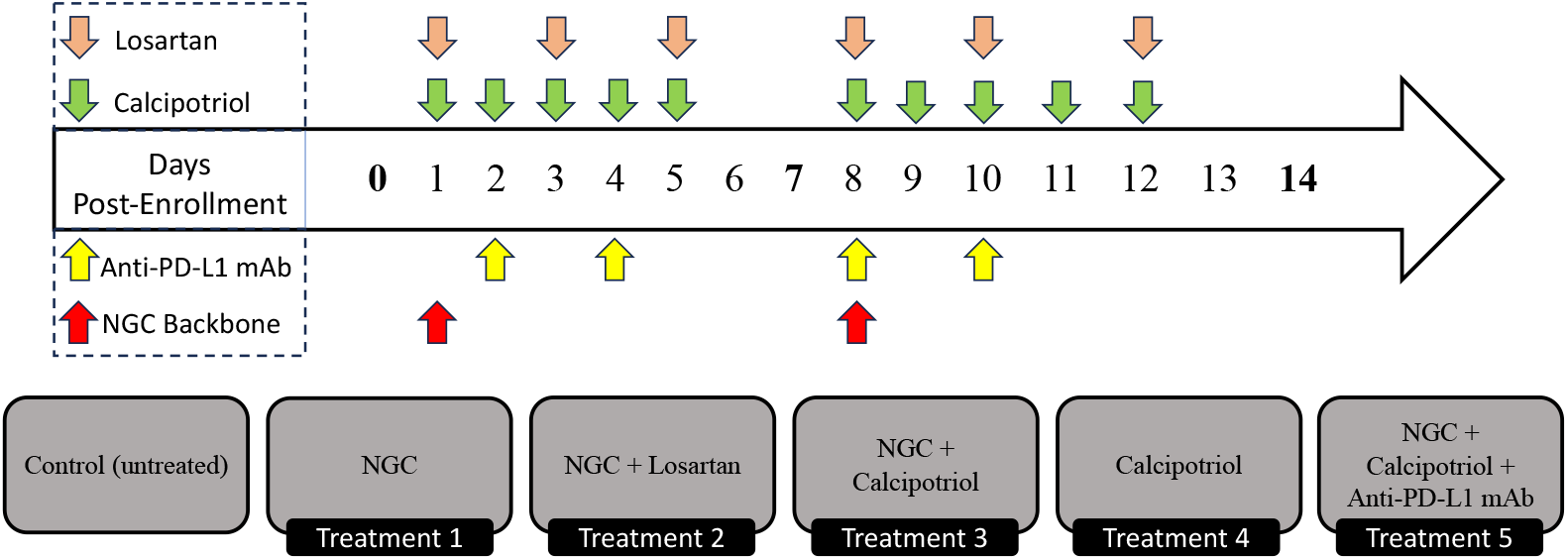
Depiction of the treatment regimes tested in the murine model of pancreatic cancer. Mice were treated for 14 days. Separate treatment components, namely NGC chemotherapy (red), anti-PD-L1 immunotherapy (yellow), and losartan (beige)/calcipotriol (green) stromal-targeting therapies, were included across five different treatment protocols (excluding control). Days with bolded time points represent tumor size measurements.

#### 2.1.3 Tumor Volume

To estimate the tumor volume, the tumor boundary was manually drawn on T2W images using the ImageJ software to generate tumor ROIs, and the areas from all ROIs was summed and the result was multiplied by the slice thickness. Tumor volume was assessed on day 0 prior to treatment, as well as on days 7 and 14 following treatment. Mice were sacrificed if a tumor measurement exceeded 1000 mm^3^.

### 2.2 Mathematical models

To characterize tumor dynamics in response to targeted treatment, we propose the calibration and prediction framework illustrated in Figure 2. We begin by defining a set of ODEs to model the temporal dynamics of pancreatic tumor volume in response to various chemo- and targeted therapies. In particular, the model accounts for tumor proliferation, drug decay rate, tumor death rate, tumor carrying capacity, and initial tumor volume. Then, we compute a sensitivity analysis to identify the most important parameters. Following this, we calibrate the models to in vivo experimental data using a Bayesian method to account for uncertainties in both the model and data. To compare the proposed models, we calculate the Bayesian Inference Criterion (BIC) and select the best model to undergo prediction analysis. Predictions are conducted in three separate scenarios: mouse-specific predictions, leave-one-out predictions, and mouse-specific group-informed predictions. To benchmark the effectiveness of these models as predictors, we evaluate their ability to successfully predict whether an individual mouse will become a treatment responder or not. We define responders as mice whose tumor volume at the end of the experimental regime is lower than their tumor volume at the start of treatment and nonresponders are mice that do not fit this criterion. In this subsection, we will derive the models, while the other steps will be presented in subsequent subsections.

**Figure 2.**
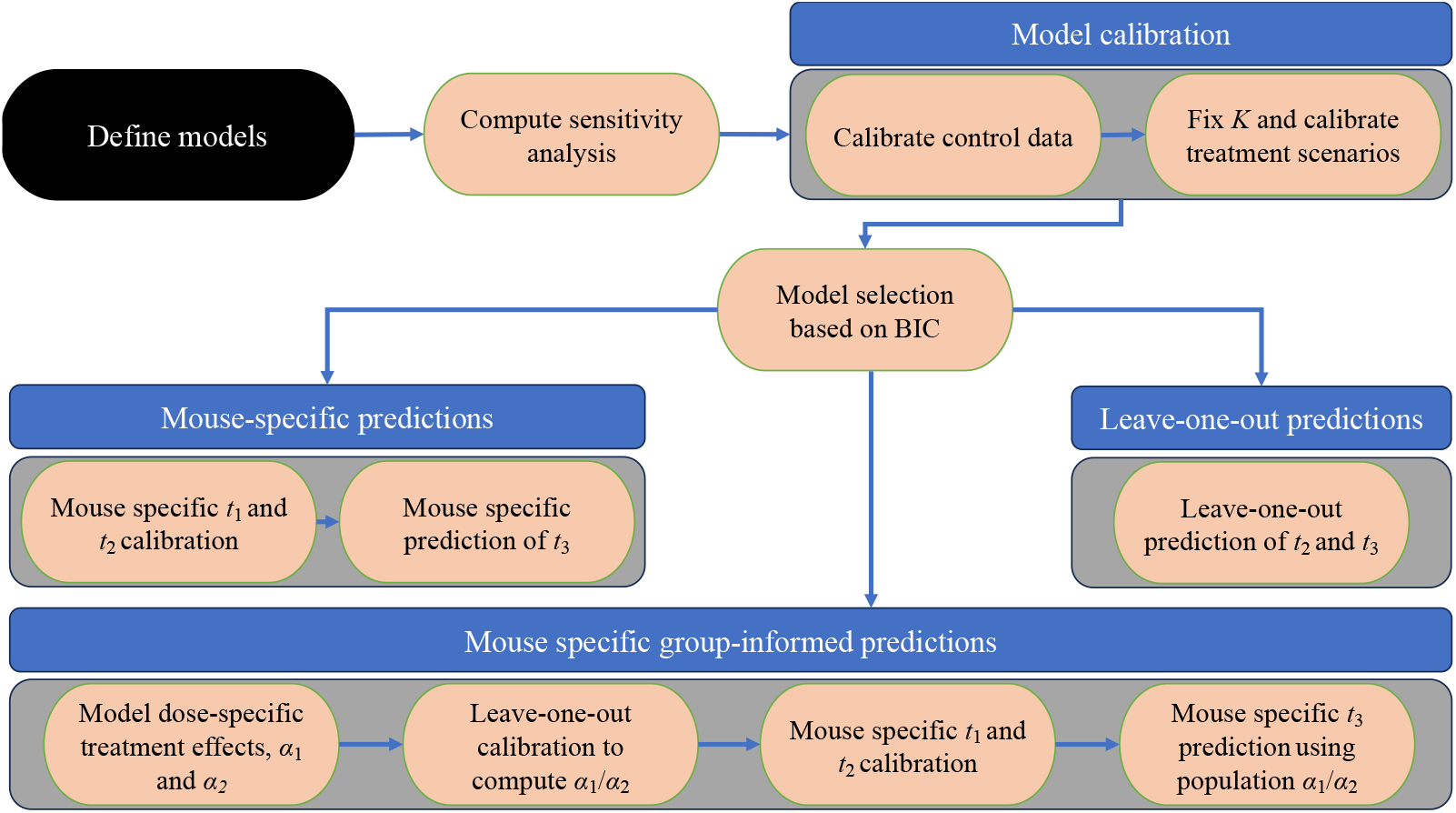
Illustration of the model calibration and prediction framework. The process begins with the definition of mathematical models designed to parsimoniously describe the treatment scenarios. Sensitivity analyses is then performed to determine the relative impact of each model parameter. After calibrating the control and treatment data to each model, the Bayesian Information Criterion (BIC) guides the selection of a single “best” model for predictions. *t*_*i*_ and *α*_*i*_ refer to the *i*^*th*^ time of tumor volume measurement and *i*^*th*^ tumor death rate due to treatment, respectively. Using the chosen model, three prediction scenarios are investigated: 1) leave-one-out predictions for days 7 and 14 tumor volumes, 2) mouse-specific calibrations using the days 0 and 7 data to predict the day 14 tumor volumes, and 3) mouse-specific, predictions made by incorporating the population’s average resistance to the treatment 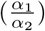 when calibrating the day 0 to day 7 data to predict day 14 tumor volumes.

In our model, we assume the tumor volume at time *t, N*(*t*) increases logistically at a proliferation rate, *r*, up to a carrying capacity of *K* (see Table 1 for a listing of all model parameters):

**Table 1.** Parameter definitions and uniform priors. *𝒰* (*a, b*) denotes a uniform distribution with bounds *a* and *b* for the prior values of the respective parameters.

| Parameter | Meaning | Prior |
| --- | --- | --- |
| $N_0$ | initial tumor volume | $\mathcal{U}(0, 600) \text{ mm}^3$ |
| $K$ | carrying capacity | $\mathcal{U}(1, 3000) \text{ mm}^3$ |
| $r$ | tumor proliferation rate | $\mathcal{U}(0, 0.3) \text{ d}^{-1}$ |
| $\alpha$ | tumor death rate by treatment | $\mathcal{U}(0, 0.3) \text{ d}^{-1}$ |
| $\beta$ | treatment effect decay | $\mathcal{U}(0, 2)$ |

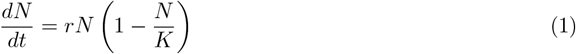

To characterize the effect of treatment in our model, we introduce a compounding linear term, with an intensity of *α*:

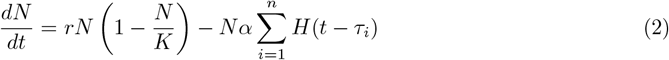

where *τ*_*i*_ is the time of the start of the *i*^*th*^ treatment interval, *α* is the death rate due to treatment, *t* is the time post initial measurement (in days), and *H* is the Heaviside function. In this treatment-agnostic model, we do not account for each specific day the treatment was delivered. Instead, we focus on the compounding effects of the treatment between key time points; specifically, between the first and second measurements (*t*_1_ and *t*_2_, respectively) and between the second and third measurements (*t*_2_ and *t*_3_, respectively). Thus, we add the effect of the treatment at *τ*_1_ = *t*_1_ and *τ*_2_ = *t*_2_. Our second treatment model extends Eq. (2) by incorporating a drug decay parameter, *β*, which allows the treatment term to exponentially decay mimicking the natural loss of drug concentration within the mice:

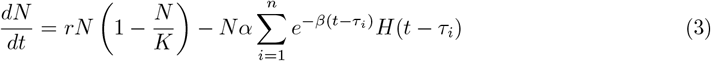

### 2.3 Sensitivity analysis

We utilize the Sobol method for performing a sensitivity analysis [27], which is a variance-based measure that determines the relative effect of independent variables on the quantity of interest (e.g., tumor volume, *N*(*t*)). We apply this method via a sampling method described by Saltelli [28, 29, 30], chosen for its efficiency in achieving convergence with a lower sample size. This application to our models is now described in detail (see Supplement 1 for visual representation). Let **M**(***θ***) represent a model defined by *Z* parameters ***θ***, which reside in the parameter space **Θ** ⊂ *ℝ*^*Z*^. We began by constructing matrices ***A*** and ***B*** by randomly sampling from a uniform distribution representing the uncertainty range of each parameter space.

Matrices ***A*** and ***B*** are of size *L × Z*, where *L* is the sample size length, and each row represents a unique set of parameters from the uncertainty space. Next, we develop *Z* matrices, 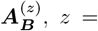, *z* = 1, 2, 3, …, *Z* and all columns are duplicated from ***A*** except the *z*^*th*^ column, which is copied from the *z*^*th*^ column of ***B***. The model is then solved for each row of matrices ***A*** and 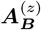 with the outputs stored in ***Y*** _***A***_ and 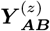, respectively, resulting in only *L* ·(*Z* + 1) model evaluations. These outputs are used to evaluate 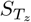, the total sensitivity index, for each parameter, *z*. We approximate the total sensitivity index for each parameter using an estimator defined by Saltelli [30]:

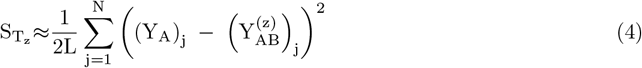

For dynamic processes (e.g., tumor growth), this form of sensitivity analysis allows us to evaluate the relative importance of individual model parameters at each time step. Thus, we can observe how the importance of each parameter changes over time to identify and eliminate unnecessary model parameters, thereby reducing model complexity. The sensitivity index is an approximation because it relies on a finite number of samples (*L*) to estimate the contribution of each parameter to the output variance. While this method provides a good estimate, the accuracy of the index improves with increasing *L*.

### 2.4 Bayesian model calibration and selection

The in vivo longitudinal experimental data described in section 2.1 is utilized to calibrate the parameters in Eqs. (1) - (3). A Bayesian framework is developed to account for uncertainty within both the data and the implemented models. The framework is defined as the following:

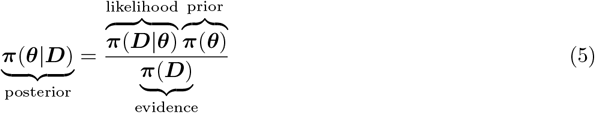

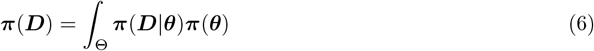

where ***D*** is the experimental data, ***θ*** is the vector of calibrated model parameters, ***π***(***θ***) is the prior estimate of a particular model parameters, ***π***(***D***|***θ***) is the likelihood that the data is observed for a set of parameters, ***π***(***D***) is the evidence (a normalizing factor), and ***π***(***θ***|***D***) is the posterior distribution of the parameters (see Table 1). We can now define a log-likelihood function as

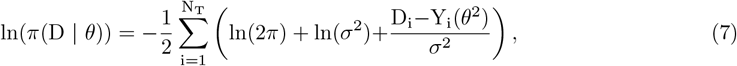

where *i* is time point and *N*_*T*_ is the number of time points. To quantitatively differentiate between the performances of the models given by Eqs. (1), (2), and (3), we calculate the Bayesian Information Criterion (BIC) [31]:

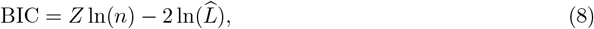

where *Z* is the number of parameters estimated, *n* is the number of observed values, and 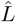 is the maximum value of the log likelihood. The model with the lowest BIC value represents the model that provides the most parsimonious description of the data, as it captures the highest likelihood of parameter value presence with a small penalty for the number of parameters, thereby prioritizing reductions in model complexity.

### 2.5 Model predictions

We develop three separate prediction scenarios to test the translational potential of our model in a clinical application. In the first scenario, we calibrate the first two time points of the available data for each mouse in each treatment group and then propagate the model solution forward to predict the tumor volume at the third and final time point for each mouse on an individual basis. The accuracy of the prediction is then compared to the experimental data via the concordance correlation coefficient (CCC). This mimics a clinical scenario in which we predict future tumor growth based on previous screenings without using heterogenous population data [32, 33, 34]. In the second scenario, we utilize a leave-one-out method and leverage population information to predict tumor volumes of individual mice in each treatment group. We began by constructing *M* matrices, ***C***_*m*_ with *m* = 1, 2, 3 … *M*, where *M* is the number of mice in a treatment group. ***C***_*m*_ is defined by the observed tumor volume for the remaining *M* − 1 mice (i.e., excluding mouse *m*) in the treatment group at each time point. ***C***_*m*_’s rows are represented by different mice in the group, whereas the columns of this matrix represent time points of tumor volume measurement. All matrices ***C***_*m*_ are then calibrated using the Bayesian framework. Then, to predict the tumor volume of a specific mouse, *m*, we randomly sample 1000 sets of parameters from the output space of the Bayesian calibration (i.e., the posterior distribution) of ***C***_*m*_ to run our forward model. Thus, this method uses a set of 1000 parameters from the treatment group and the initial tumor volume to characterize a Bayesian distribution for future tumor volumes; namely, the tumor volumes at *t*_2_ and *t*_3_, where *t*_*i*_ represents the time of the *i*^*th*^ tumor volume measurement.

For scenario 3, we develop a hybrid of the mouse-specific and leave-one-out predictions. We began by splitting the drug effect into two separate linear summation terms, each representing the individual treatment doses administered on different days. This approach allows us to introduce a new correction factor for tumor-specific parameters and to account for the possibility that the tumor’s response to treatment may change after the first dose. For example, even if the doses are the same on both treatment days, the effect of the treatment might differ due to factors such as the tumor developing some level of resistance or other changes in its response over time. To account for this phenomenon, we modify Eq. (2) as follows:

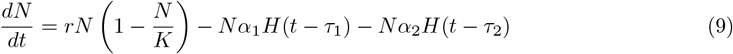

where *α*_1_ is the death rate due to treatment received between times *t*_1_ and *t*_2_, and *α*_2_ is the death rate due to the treatment received between times *t*_2_ and *t*_3_. As described in Figure 1, we have five treatment combinations, and this same model is used for each treatment. Therefore, the parameters *α*_1_ and *α*_2_ can represent different treatments and combinations, with their values being calibrated to match each specific treatment protocol. This allows us to develop relationships between treatment administration intervals. To account for the effect of serial doses, we make the simplifying assumption that *α*_2_ is proportional to *α*_1_, meaning we can describe *α*_2_ as a linear function of *α*_1_. The proportionality between *α*_2_ and *α*_1_, which we refer to as treatment resistance, is computed as the average ratio of *α*_1_ to *α*_2_ across the group of mice. To implement this concept of treatment resistance and predict the final tumor volume for mouse m, we begin by calibrating the model given by Eq. (9) to experimental data from every other mouse in the treatment group (i.e., *M* − 1), thereby determining the values of *α*_1_ and *α*_2_ for those mice. For mouse *m*, we then calibrate only *α*_1_ and use the treatment resistance ratio from the group to estimate *α*_2_ (which has not been directly calibrated for mouse *m*). This group-derived proportionality is applied to adjust *α*_2_ before continuing to forward propagate the model. Thus, for a single mouse, we combine our mouse-specific calibration with the group-derived proportionality of compounding treatments to derive a prediction for the final tumor volume. Through this method, we effectively combine known group information regarding the effect of serial dosage while still maintaining patient-specific modeling of tumors to create predictions.

### 2.6 Numerical implementation

Eqs. (1) - (3) are implemented in Python 3.11.9 and the calibration framework is illustrated in Figure 2. The computation of the posterior density is performed through a parallel, adaptive, multilevel Markov Chain Monte Carlo sampling technique, available through the emcee Python library [35]. The ODEs are solved using a fourth-order Runge-Kutta method. Detailed information on the code, including instructions on how to run it and the necessary dependencies, is available at https://github.com/krithikvishwanath.

### 2.7 Statistics and reproducibility

Statistical analysis between Bayesian treatment groups is calculated using a two-tailed Mann-Whitney U test on tumor volumes. To do so, we note that groups are independent of each other per experimental protocol, and tumor volume is an ordinal quantity. Bayesian groups each contain 10000 samples to represent the distribution. To evaluate the effects of growth between initial and final tumor volumes of a particular group (i.e., to demonstrate that the tumor is indeed growing in the control), we utilize a paired, two-tailed t-test. To assess statistical significance, a Bonferroni-adjusted *p*-value was used to maintain a 5% probability of a Type-I error.

## 3 Results

### 3.1 Sensitivity analysis

Utilizing the parameter ranges detailed in Table 1, we performed temporal sensitivity analysis on both the linear (Eq. (2)) and the exponential decaying (Eq. (3)) treatment models. Figure 3 presents the results covering the same temporal period as the experimental regime, spanning days 0 through 14. The sensitivity analysis for both models reveals that at lower time indices (i.e., before days 7 and 9 for the linear and exponential decaying treatment models, respectively), the initial tumor volume, *N*_0_, is the most influential parameter. This suggests that small changes in this value would result in notably different tumor volumes during this time interval. However, by the end of the experiment (i.e., day 14), the initial tumor volume becomes the least important parameter in determining the final tumor volume. For the linear model, the final tumor volume is primarily influenced by the death rate due to treatment, *α*, followed by the proliferation rate of the tumor, *r*, and the carrying capacity, *K*, with *N*_0_ becoming the least influential parameter. Conversely, for the exponentially decaying model, the most important parameter is the carrying capacity, *K*, closely followed by the initial proliferation rate, *r*. In this model, both the death rate due to treatment, *α*, and the decay rate of the treatment effect, *β*, have a negligible effect on the tumor volume in comparison to the other parameters with a total sensitivity index, a dynamic measure of parameter influence on an output variable (i.e., tumor volume), consistently lower than 0.1 throughout the entire time interval. These results suggest that in the linear model, the death rate due to treatment is the most crucial parameter influencing tumor dynamics. In contrast, in the exponential decaying model, the decay term diminishes the importance of the treatment during the observed interval, making the carrying capacity and the proliferation rate the parameters that need to be precisely calibrated.

**Figure 3.**
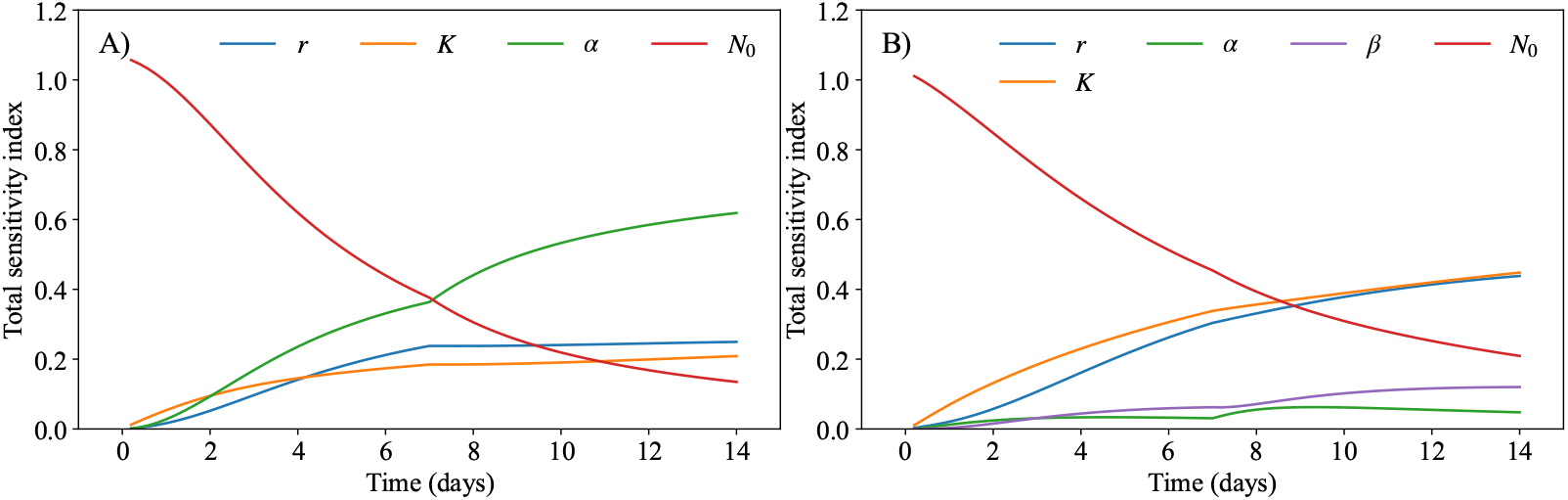
Sensitivity analysis of the logistic model with a linear treatment term (Eq. (2); panel A) and the logistic model with a decaying treatment term (Eq. (3); panel B). The total sensitivity index (Eq. (4)) is computed for each model, demonstrating how the parameters affect tumor volume throughout the experimental time course. In both panels, the initial tumor volume plays a large role before the second dose of treatment (day 7), but its effect reduces as the experiment continues. In panel A, the death rate due to treatment (*α*, green line) has the highest total sensitivity index by far (approximately 3× higher than the second most influential parameter), followed by the proliferation rate. However, for panel B, the carrying capacity (*K*, orange line) and proliferation rate (*r*, blue line) exhibit nearly identical total sensitivity indices by day 14, with similar temporal dynamics throughout the experiment. Both parameters emerge as the most influential factors after day 9. These parameters are followed in total sensitivity index by the initial tumor volume (*N*_0_, red line). This discrepancy between panel A and panel B’s high-order parameters is likely due to the lack of decay present for the treatment in the panel A model, resulting in a larger value of the death rate due to treatment parameter, *α*.

### 3.2 Calibration of the control group to the logistic model

Continuing the framework outlined in Figure 2, we employ the priors defined in Table 1 to calibrate the parameters *r, K*, and *N*_0_ to the control data. In this scenario, we calibrate a population-specific *K* (i.e., one distribution of the parameter that fits the entire group), along with a mouse-specific *r* and *N*_0_ (i.e., each mouse having its own distribution of *r* and *N*_0_). As depicted in Figure 4, the logistic model (i.e., Eq. (1)) can accurately describe the experimental data from the control group, with a CCC and PCC of 0.99 when comparing the experimental and calibrated tumor volumes, with a mean absolute percent error (MAPE) below 8%. Moving forward, we use these parameters when calibrating the models described by Eqs. (2) and (3). Specifically, the prior distribution for the Bayesian calibration of *r* for each mouse in the treatment groups is established using the upper and lower bounds determined from the posterior distribution of the control group’s proliferation rate. Additionally, the carrying capacity calculated from the control group, *K*, is set to the maximum likelihood value from this calibration when calibrating the treated groups. This allows us to reduce the number of parameters requiring calibration in the other models and mitigate potential issues with parameter identifiability.

**Figure 4.**
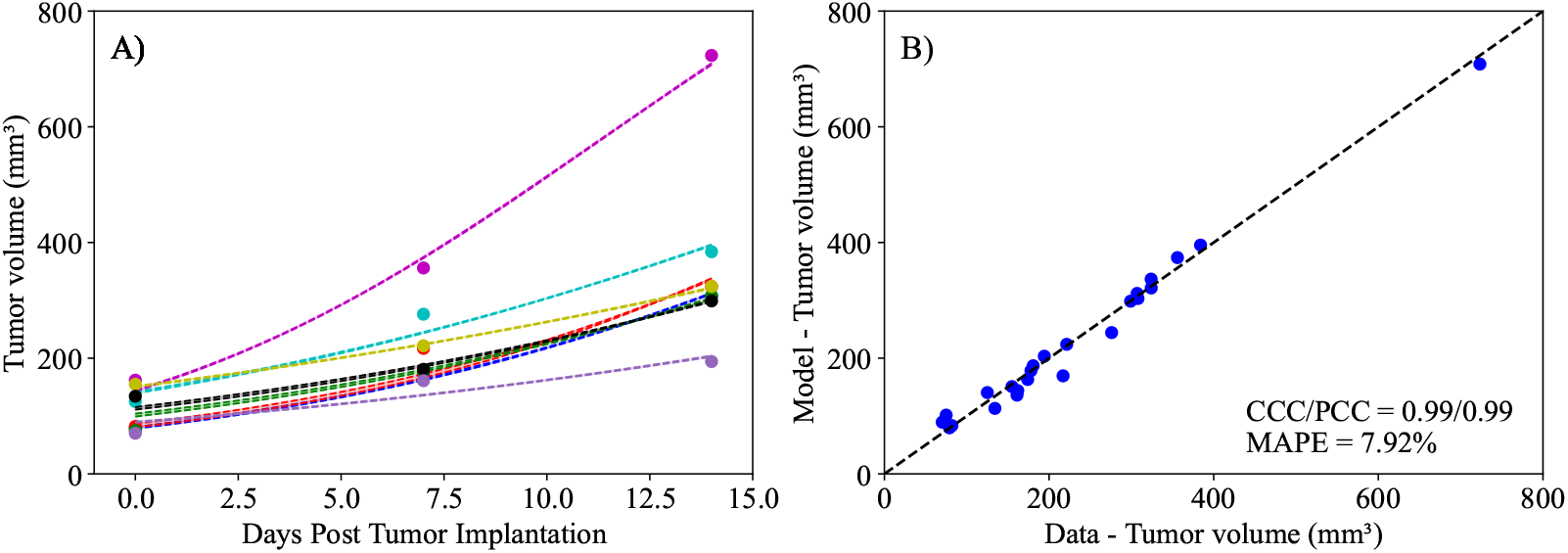
Calibration of control data to the logistic model described by Eq. (1). Panel A presents calibration of the model to individual mouse tumor volumes (solid points) at three time points (days 0, 7, and 14) over the two-week experimental period, along with their corresponding calibrated curve (dashed lines). Panel B compares the experimentally measured tumor volume and the model’s calibrated maximum likelihood value. The dashed black line in Panel B is the line of unity. The calibrated logistic model demonstrates exceptional alignment with the control data, achieving a concordance correlation coefficient (CCC) and a Pearson correlation coefficient (PCC) of 0.99 each, along with a mean absolute percent error (MAPE) of 7.92%. As the control group received no treatment, by day 14 all mice exhibit a significantly higher final tumor volume compared to their initial tumor volume on day 0 (two-tailed paired t-test, *p* = 0.0013).

### 3.3 Calibration of treatment-based models

Following the calibration of Eq. (1) to the control data, we proceed to calibrate the parameters in the linear (Eq. (2)) and the exponential decaying (Eq. (3)) treatment models to the data from each treatment protocol. In both models, we calibrate the parameters *r, α*, and *N*_0_ for each mouse, while fixing the carrying capacity to the value obtained from fitting the control data to Eq. (1). Additionally, Eq. (3) requires the calibration of *β* (i.e., the treatment decay effect) as a populationbased parameter, with each treatment protocol having its own distribution of *β*. Notably, the models in consideration are treatment agnostic and do not depend on the specific nature of any individual treatment combination. In Figure 5, we present the experimental data and the model solutions for all five treatment scenarios. The model differentiates between responders and non-responders with 100% accuracy in all treatments except Treatment 1, where the model achieved a 94.44% accuracy. The mean absolute percent error for the tumor volume is below 10%, and the CCC/PCC is above 0.98 for all treatment protocols. Similarly, we display the results for the second model (i.e., Eq (3)) in Figure 6. This model can differentiate between responders and non-responders with an accuracy of 100% in all scenarios except treatment protocols 1 and 5, where the model achieved an accuracy of 94.44% and 87.50%, respectively. Notably, the only discernible difference in terms of accuracy between the two treatment-agnostic models occurs in Treatment 5, where the second model (i.e., Eq. (3)) incorrectly classifies one non-responder as a responder. The CCC/PCC for the second model is above 0.95 for all treatment protocols, and the mean absolute percent error for tumor volume is below 10% for all treatment scenarios. These results indicate that both models effectively capture the effects of different tumor protocols on the tumor volume.

**Figure 5.**
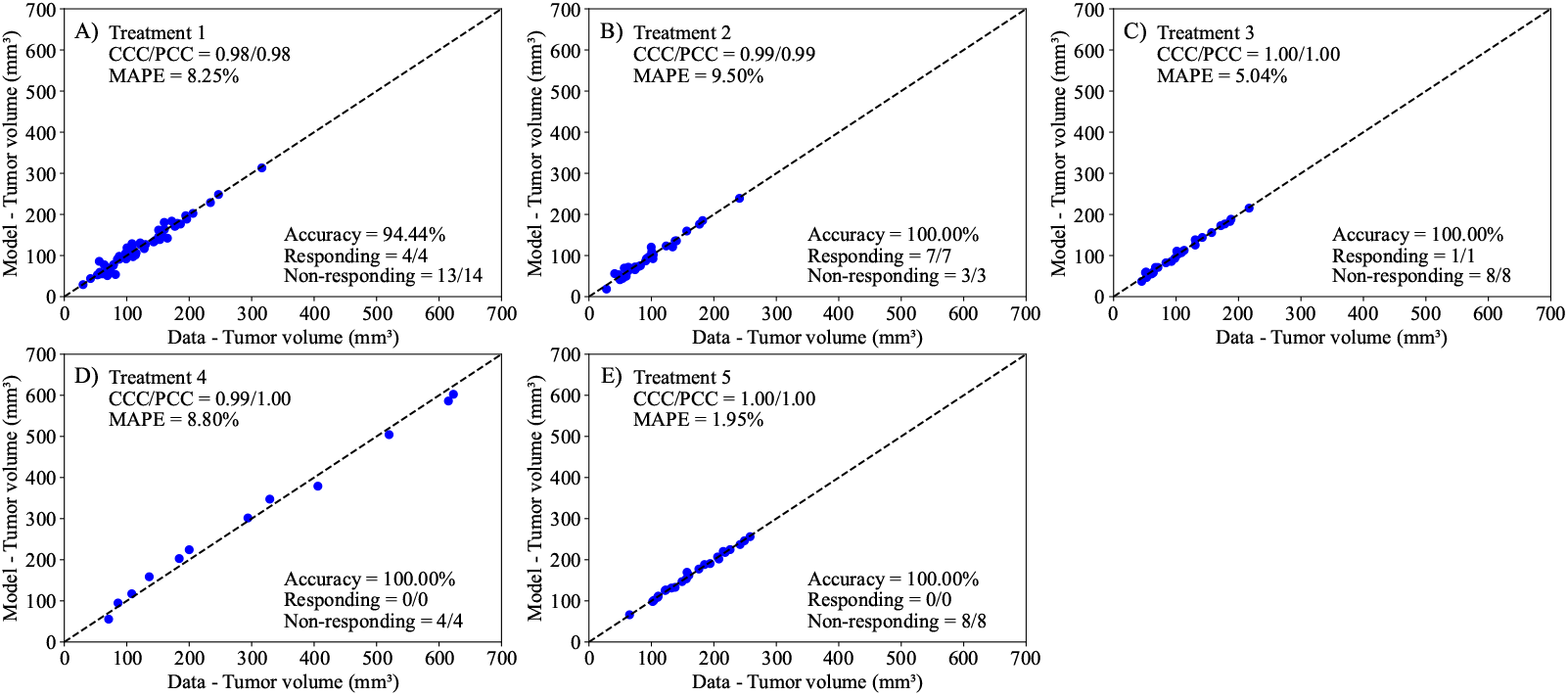
Comparison between the experimentally measured tumor volume and model calibration of logistic growth with a linear treatment term (Eq. (2)) for each treatment protocol. The dashed black lines are the line of unity. All calibrations to treatment scenarios exhibit high levels of correlation between the data and model, with all CCCs and PCCs greater than 0.98. The average accuracy, calculated as the number of correctly identified responders and non-responders divided by the total number of mice for each treatment, was 98.89 ± 1.11% across all treatment scenarios. Further, the MAPE for all scenarios is less than 10%. This suggests that the Bayesian calibration of model parameters (*r, α*, and *N*_0_) effectively reproduces the experimental data.

**Figure 6.**
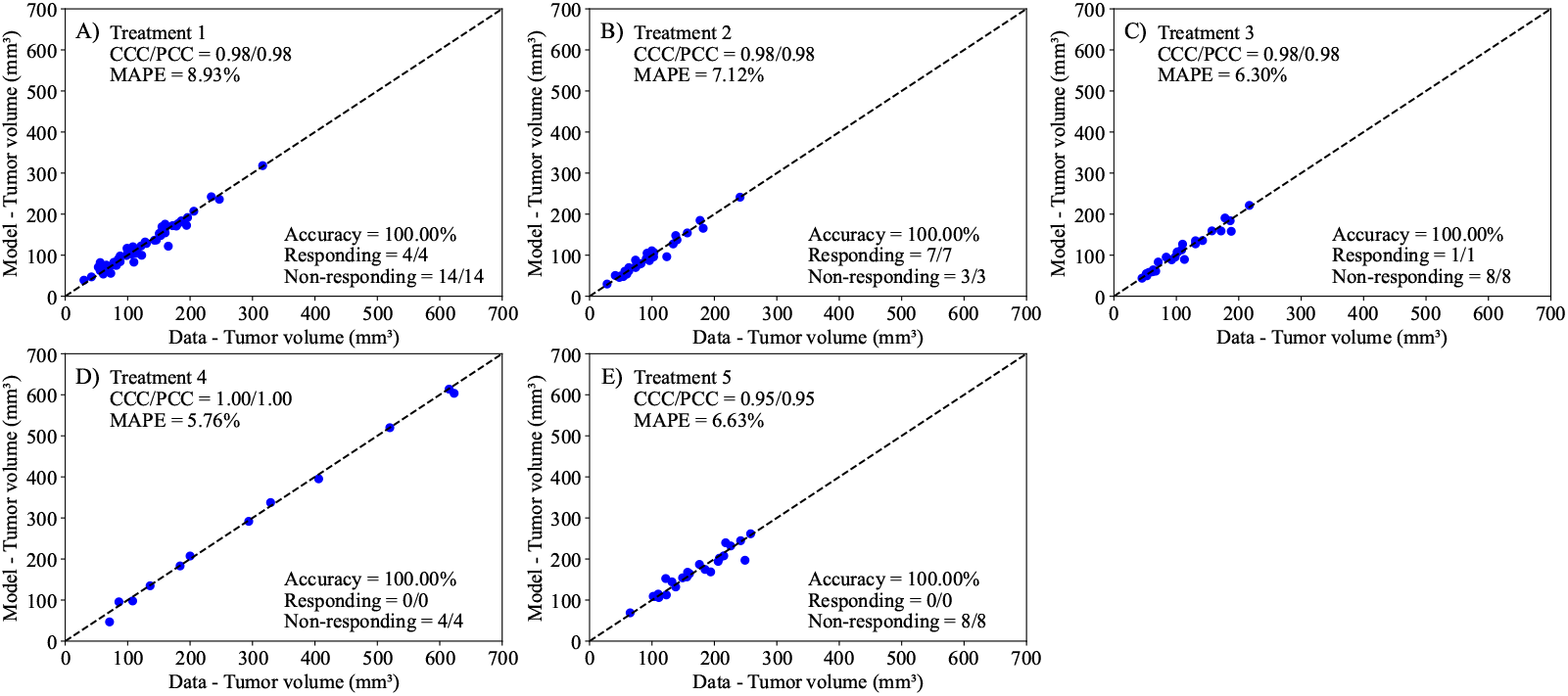
Comparison between the experimentally measured tumor volume and model calibration of logistic growth with a drug decay treatment term (Eq. (3)) for each treatment protocol. The dashed black lines are the line of unity. All calibrations to treatment scenarios exhibit high levels of correlation between the data and model, with all CCCs and PCCs greater than 0.95. The average accuracy was 100% across all treatment scenarios. Further, the MAPE for all scenarios is less than 10%. This suggests that the Bayesian calibration of model parameters (*r, α, β*, and *N*_0_) effectively reproduces the experimental data.

### 3.4 Model selection

In Table 2, we present the Bayesian Information Criterion (BIC) for each model (linear and exponentially decaying) for each treatment protocol. Note that a lower BIC indicates a better fit of the model to the data [31]. Our results show that the linear model, Eq. (2), consistently performs better in every treatment protocol and has an average BIC of 46 lower than the exponentially decaying model, Eq. (3). Therefore, this model is selected for the subsequent analyses in this study.

**Table 2.**
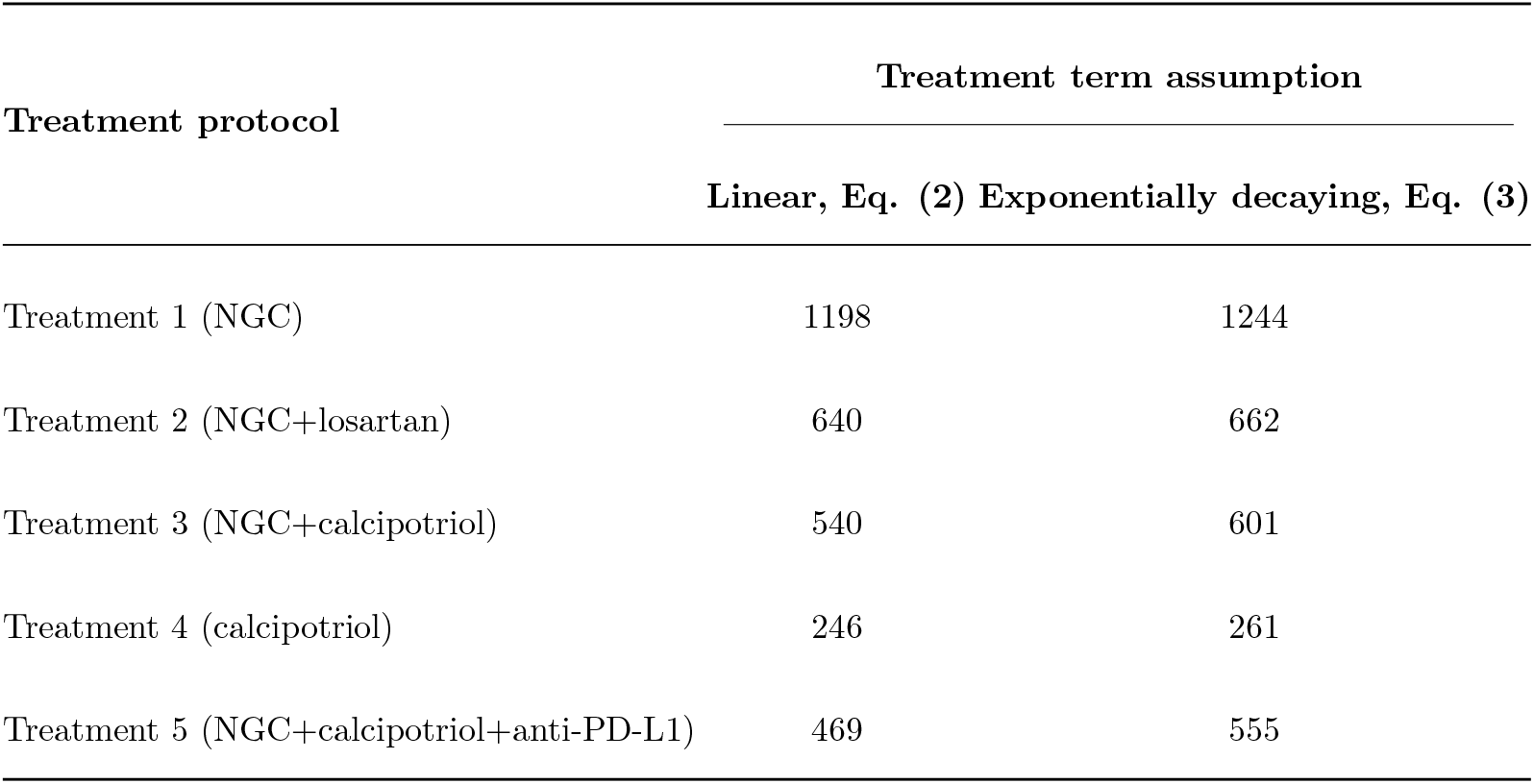
Calculated Bayesian Inference Criterion (BIC) on model calibration for parameters in Eq. (2) and Eq. (3), based on maximum likelihood and a parameter cardinality penalty. See Figure 1 for the list of the drugs included within each treatment protocol.

| Treatment protocol | Treatment term assumption |  |
| --- | --- | --- |
|  | Linear, Eq. (2) | Exponentially decaying, Eq. (3) |
| Treatment 1 (NGC) | 1198 | 1244 |
| Treatment 2 (NGC+losartan) | 640 | 662 |
| Treatment 3 (NGC+calcipotriol) | 540 | 601 |
| Treatment 4 (calcipotriol) | 246 | 261 |
| Treatment 5 (NGC+calcipotriol+anti-PD-L1) | 469 | 555 |

### 3.5 Parameter distributions of model calibration

After selecting the model with the linear treatment term defined by Eq. (2) as the best model to represent the data (according to the BIC), we proceed to examine the posterior parameter distributions for each of the control and treatment protocols. Figure 7 presents significant differences across all scenarios for the tumor proliferation rates, *r*, and tumor death rate due to treatment, *α*. In Panel A), the median proliferation rate for non-responders is significantly higher than that for responders across Treatments 1, 2, and 3 (23.44 ± 14.50% higher than responders, *p*-adjusted *<* 0.001). Treatment 4 has the highest proliferation rate, while Treatments 2 and 3 have the lowest. Except for the responders in Treatment 2 and 3, all treatment groups have a median proliferation rate that significantly surpasses the control group. Panel B) displays the distribution of the parameter for the death rate due to treatment. In scenarios with both responders and non-responders (i.e., Treatments 1-3), the median death rate of non-responders is 54.8 ± 5.54% lower (*p*-adjusted *<* 0.001) than that of responders. Statistical analysis using a Bonferroni-adjusted Mann-Whitney U test indicates that differences between all pairwise groups (i.e., responders and non-responders) in terms of tumor proliferation rates and tumor death rates due to treatment are significant (*p*-adjusted *<* 0.001). The lowest death rate due to treatment is associated with Treatment 4, while the highest is observed in the responders in Treatment 1.

**Figure 7.**
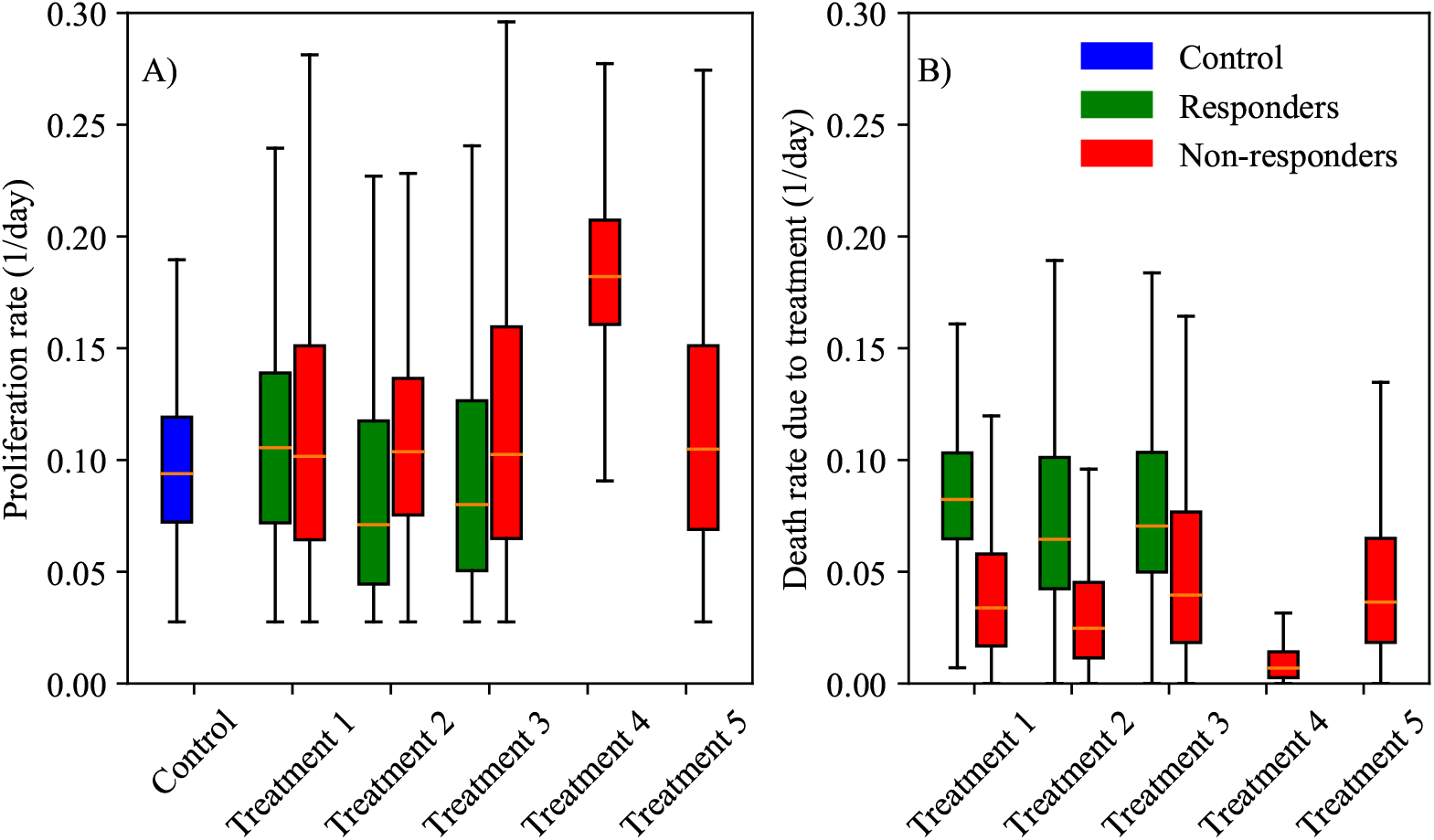
Box and whisker plots of posterior parameter distributions for each treatment protocol. Specifically, parameter distributions for proliferation rate (*r*) and death rate due to treatment (*α*) are displayed in panels A and B, respectively. Bayesian distributions are displayed for control (blue) and each treatment, split into responders (green) and non-responders (red). Responders exhibit statistically lower proliferation rates and greater death rates due to treatments (*p*-adjusted *<* 0.001). In particular, Treatment 2 and Treatment 3 induce the greatest death rate due to the regimes. Treatment 4 performed the worst, with both the highest tumor proliferation rate and lowest death rate due to treatment. Note that neither Treatment 4 nor Treatment 5 had any responders.

### 3.6 Calibration of model with cumulative drug effect

In this calibration, our goal is to quantify the compounding effect of the treatment regime by calibrating a different *α* at each of the two intervals of treatment administration, effectively separating the death rate due to treatment into *α*_1_ and *α*_2_ (corresponding to the death rates caused by the first and second administration intervals of treatment, respectively). We calibrate parameters *r, α*_1_, *α*_2_, and *N*_0_ to the experimental data, and the results are presented in Figure 8. This modified treatment term of the model in Eq. (2) successfully captures the experimental data, with a CCC/PCC greater than 0.96 for all treatment scenarios. Further, the model differentiates between responders and non-responders with a 100% accuracy in all scenarios except Treatment 1, where the accuracy is 83.33%. Due to the presence of only two treatment administration intervals in each protocol, we assume the compounding effect occurs in a linear fashion; that is, we model each treatment as directly proportional to the previous treatment administration interval by a constant, defined as treatment resistivity 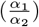. According to this definition, a higher treatment resistivity indicates a decrease in impact for the second dose of treatment relative to the first. Our findings reveal that across all treatment protocols, the median resistivity to treatment is between 0.72 and 4.50, indicating that the first delivery of treatment is between 0.72 and 4.5 times more effective than the second delivery of treatment. (Note that any value above 1.0 indicates the second treatment was less effective than the first.) Figure 9 presents the posterior distribution of 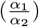 for each group. Of all treatment protocols, the responders and non-responders in treatment protocol 1 have significantly lowest resistivity to treatment (median 0.72 and 1.26, respectively; *p*-adjusted *<* 0.001), while responders in Treatment 3 and non-responders in Treatment 4 exhibit the highest resistance to treatment (4.50 and 3.2, respectively; *p*-adjusted *<* 0.001).

**Figure 8.**
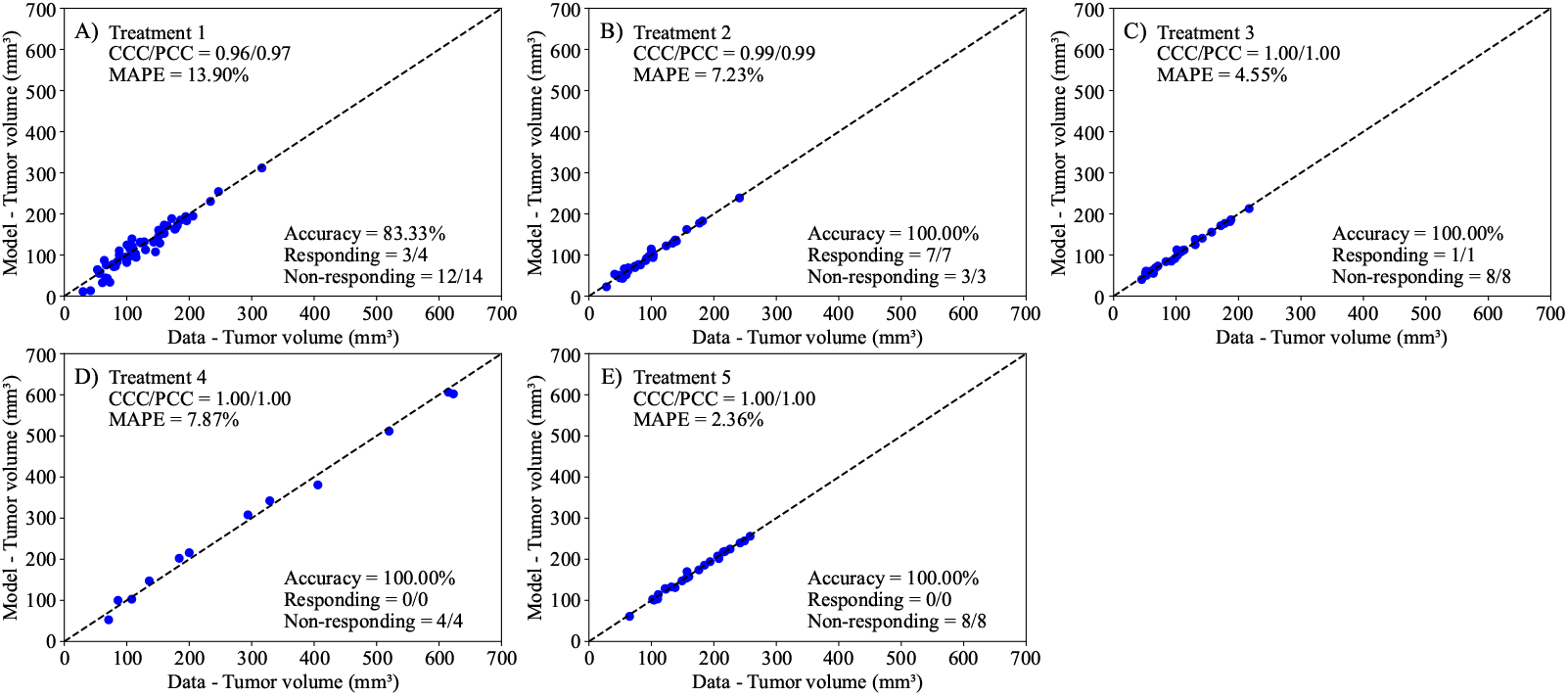
Comparison between experimentally measured tumor volume and model calibration of logistic growth with two linear treatment terms (Eq. (2)). The model now includes the effects from two separate treatments death rates (*α*_1_ and *α*_2_) to account for the individual effects of both treatment days. The dashed black lines are the line of unity. All treatment scenarios exhibit high levels of correlation between the data and the model, with all CCCs and PCCs greater than 0.96. The average accuracy was 96.67 ± 3.33% across all treatment scenarios. Further, the MAPE for all scenarios is less than 14%. This suggests that the Bayesian calibration of model parameters (*r, α*_1_, *α*_2_, *β*, and *N*_0_) effectively reproduces the experimental data.

**Figure 9.**
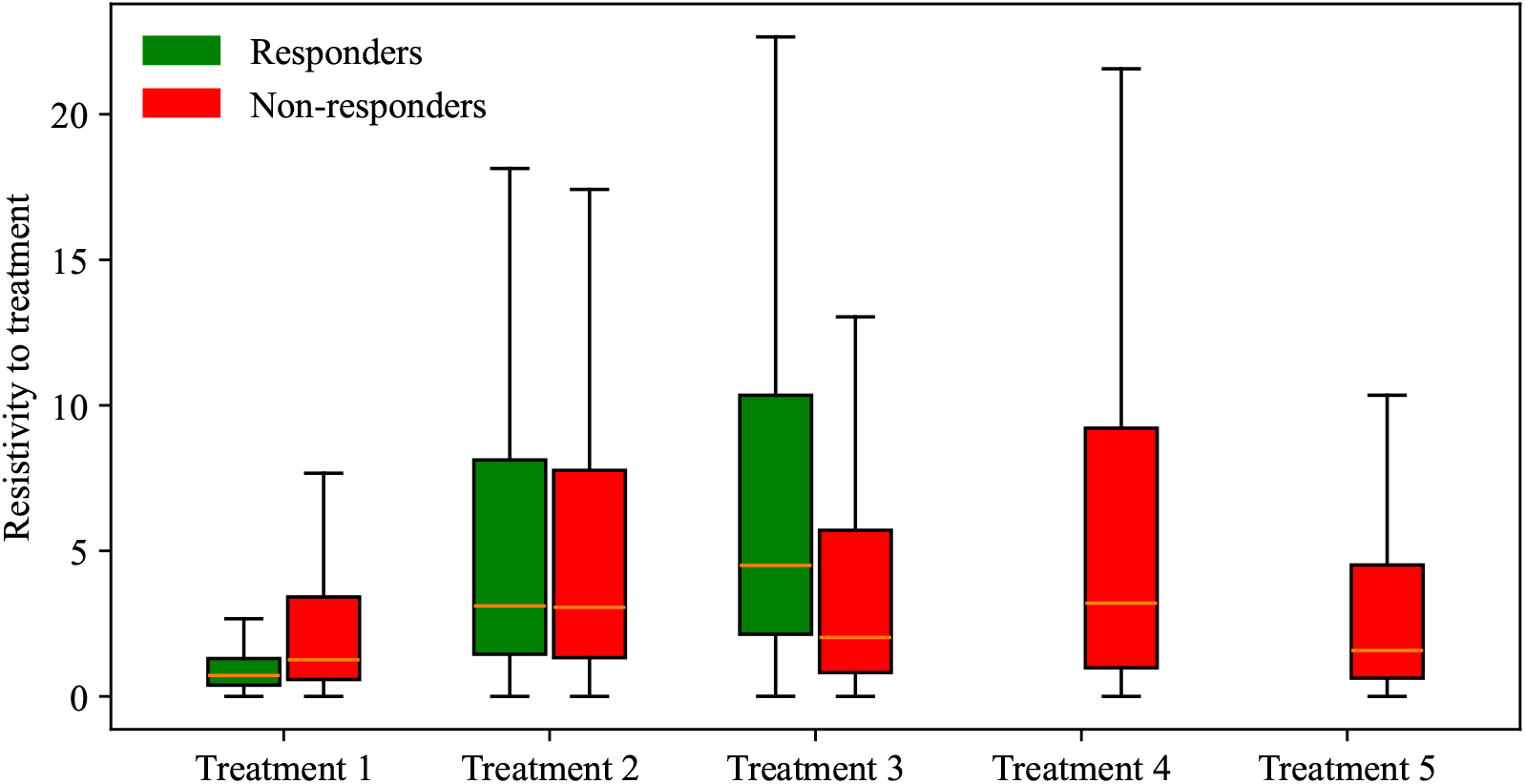
Box and whisker plots indicating the range of resistivity values encountered across distinct treatment scenarios. Each treatment scenario’s posterior distribution for resistivity to treatment, split into responders (green) and non-responders (red), is presented. The magnitude of treatment resistivity represents the ratio of the effect of the first dose (*α*_1_) to the effect of the second dose (*α*_2_); i.e., 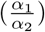. For all groups, the interquartile range is between 1 and 10, indicating a significantly lower efficacy of the second dose compared to the first. Statistical analysis using the Bonferroni-adjusted Mann-Whitney U test shows that the differences in treatment resistivity between groups are significant (*p*-adjusted *<* 0.001). Out of all scenarios, Treatment 1 has the lowest resistivity to treatment.

### 3.7 Model predictions

Following the framework presented in Figure 2, the next step is to predict tumor volumes using our selected model. We apply three prediction scenarios: leave-one-out predictions (Figure 10), mouse-specific predictions (Figure 11), and group-informed, mouse-specific predictions (Figure 12). By utilizing only the first time point and the group-informed parameter samples, the model can differentiate between responders and non-responders with an average accuracy 72.83 ± 16.10% across all treatment protocols. Additionally, for all cases except for Treatment 2 (NGC Backbone with Losartan), both the CCC and PCC are above 0.7. The MAPE remains below 33% across all scenarios, with the smallest percent error observed in Treatment 5 (MAPE = 14.75%) and the highest in Treatment 2 (MAPE = 32.22%).

**Figure 10.**
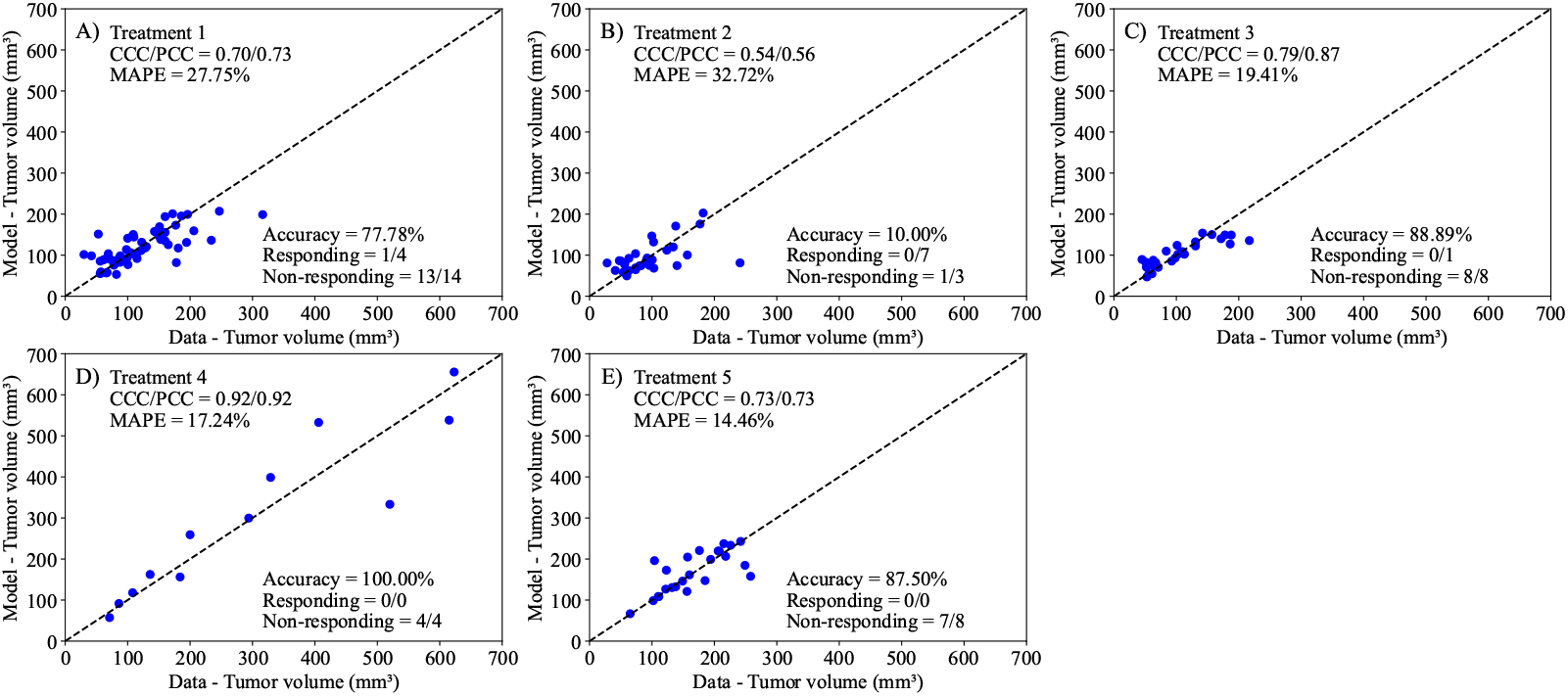
Comparison of tumor volume from experimental data and tumor volume from leave-one-out predictions for each treatment scenario. The dashed black lines are the line of unity. The average accuracy was 72.83 ± 16.10% across all treatment scenarios. Notably, leave-one-predictions exhibited greater than 75% accuracy of correctly differentiating between responders and non-responders, and a CCC and PCC greater than 0.7 for all scenarios other than Treatment 2. The MAPE is below 33% for all scenarios, performing with the lowest percent error for Treatment 5 (MAPE = 14.75%) and the greatest for Treatment 2 (MAPE = 32.22%).

**Figure 11.**
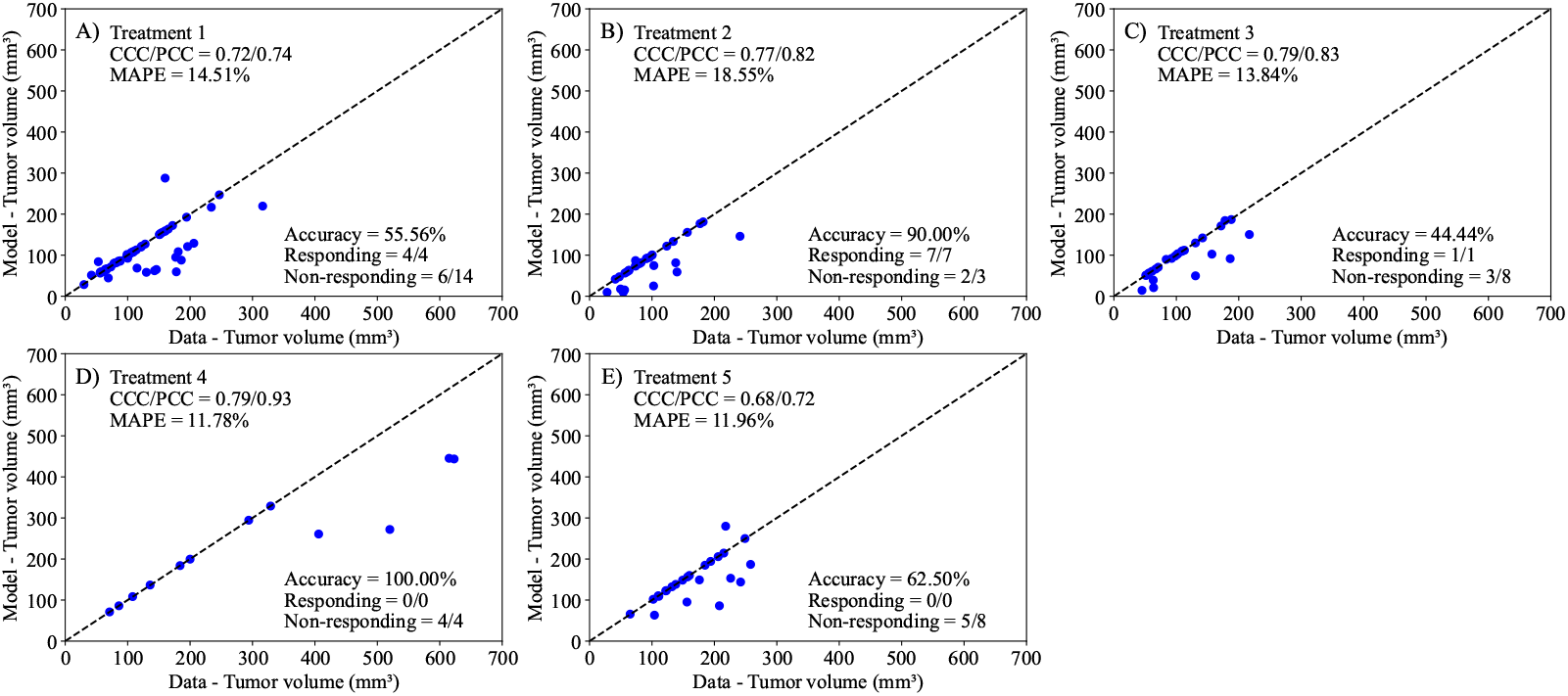
Comparison of tumor volume from experimental data and tumor volume from mousespecific predictions for each treatment scenario. The dashed black lines are the line of unity. The average accuracy was 70.50 ± 10.53% across all treatment scenarios. Mouse-specific predictions exhibited a concordance correlation coefficient (CCC) and Pearson correlation coefficient (PCC) greater than 0.68 for all scenarios. The mean absolute percent error (MAPE) is below 18.54% for all scenarios, performing with the lowest mean percent error for Treatment 5 (MAPE = 11.96%) and the greatest mean percent error for Treatment 2 (MAPE = 18.55%).

**Figure 12.**
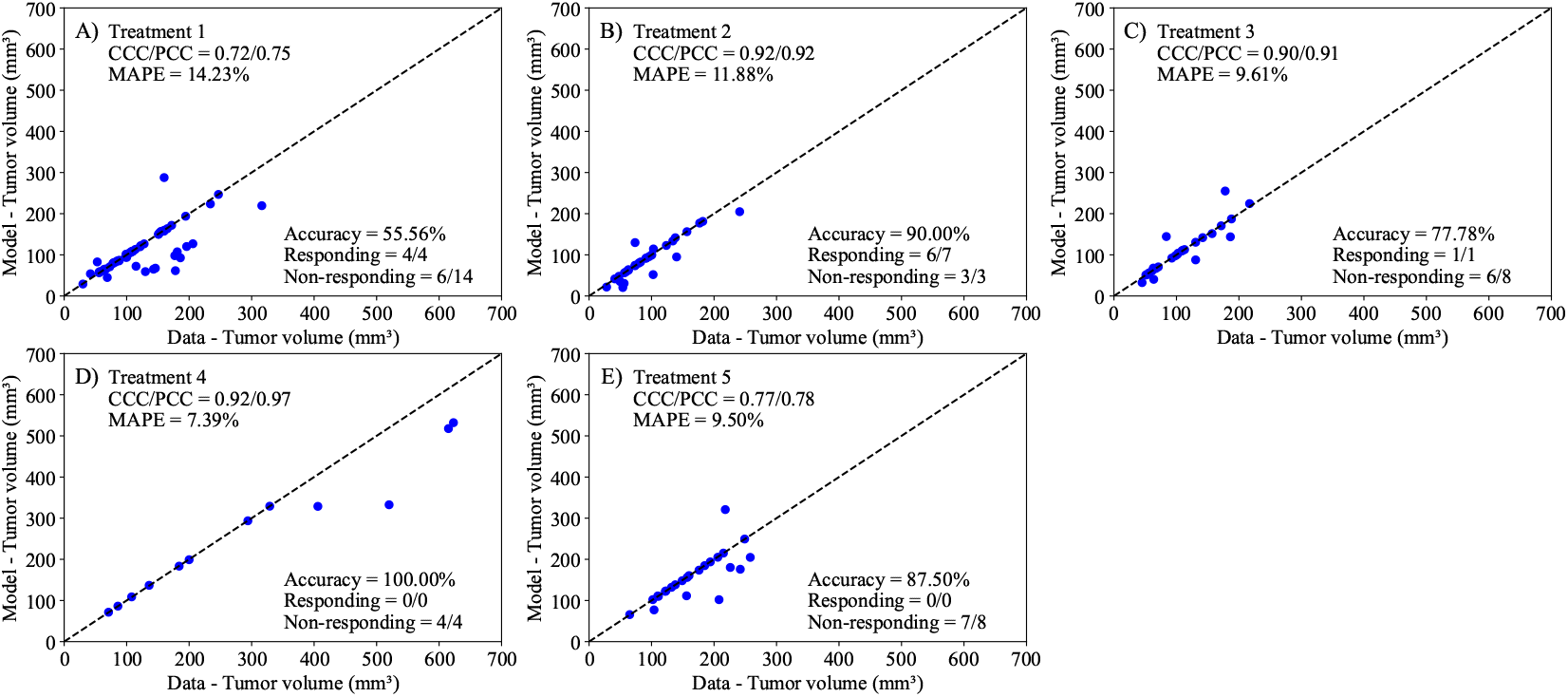
Comparison of tumor volume from experimental data and tumor volume from groupinformed, mouse-specific predictions for each treatment scenario. Calibrations utilize the population resistivity dynamics to predict tumor volumes for specific mice within the population. The dashed black lines are the line of unity. The average accuracy was 82.17 ± 7.53% across all treatment scenarios. Group-informed, mouse-specific predictions exhibited a CCC and PCC greater than 0.72 in all scenarios. The MAPE is below 14.32% for all scenarios, performing with the lowest mean percent error for Treatment 4 (MAPE = 7.29%) and the greatest mean percent error for Treatment 1 (MAPE = 14.32%).

The second prediction scheme involves mouse-specific predictions, where the parameters *r, α*, and *N*_0_ are calibrated to each mouse using the data from days 0 and 7. Following this calibration, we predict the tumor volume at day 14. In Figure 11, we compare the model predictions to the true experimental data on day 14. The model successfully differentiates responders and non-responders 70.50 ± 10.53% across all treatments, with both the CCC and PCC surpassing 0.68 for every treatment condition. The MAPE remained under 18.55% for each scenario, showing the best precision for Treatment 5 with a MAPE of 11.96%, while Treatment 2 had the highest error at 18.55%. Overall, the model underpredicts the final mice volume and had higher rates of successfully predicting responders (12/12) rather than non-responders (20/37). This observation may be attributed to the model being calibrated only for the first 7 days, potentially indicating differences in treatment efficacy during the second period of the treatment regimen (from day 7 to day 14).

Our final prediction scenario, the group-based, mouse-specific prediction method, builds upon the mouse-specific prediction by incorporating into the model the diminishing effectiveness of repeated treatments, as illustrated in Figure 9. Similar to the first prediction scheme, we calculate groupaveraged resistance to treatment 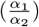 within each treatment protocol. We use this value to adjust the posterior distribution of the individual mouse’s *α*_1_ and estimate *α*_2_. This is then integrated as the new predicted death rate of our second treatment dose to predict treatment effects during the second day of treatment. Similar to our mouse-specific prediction scheme, the parameters are calibrated using the tumor volume data from days 0 to 7, and the model then predicts the final tumor volume on day 14. The outcomes of our group-based, mouse-specific predictions are presented in Figure 12. This integrated method successfully distinguishes between treatment responders and non-responders, achieving an average accuracy of 82.17 ± 7.53% across all treatment scenarios. Moreover, the prediction model attains a CCC/PCC of over 0.9 for Treatments 2, 3, and 4, accurately reflecting the actual experimental outcomes for these groups. For Treatments 1 and 5, the model achieves a CCC/PCC of over 0.72. Lastly, the MAPE remains below 15% for all scenarios, with the best performance observed in Treatment 4 (7.29%) and the least favorable in Treatment 1 (14.32%).

## 4 Discussion

We developed two mathematical models, one with linear treatment effects and one with exponentially decaying treatment effects, to characterize and predict the response of pancreatic tumors to 6 combinations of chemotherapy, stromal-targeting drugs, and immunotherapy. As both models are identical in the absence of treatment (representing logistic tumor growth), we first calibrated the control group, achieving a CCC of 0.99 (Figure 4), and used the carrying capacity value obtained here when calibrating the two models. The calibration of the linear model provided a CCC of 0.99 ± 0.01 across the five treatment groups (Figure 5), while the exponentially decaying model resulted in a CCC of 0.98 ± 0.02 (Figure 6). After calibration, the BIC selected the linear treatment model as the most parsimonious (Table 2). This model demonstrated a high degree of accuracy in fitting the in vivo data, effectively capturing the complex interactions between tumor cells, stromal components, and immune cells, allowing for robust predictions of tumor growth and regression. According to the results from the sensitivity analysis (Figure 3), in the model with exponentially decaying treatment effects, both the death rate due to treatment, *α*, and the decay rate of the treatment effect, *β*, have a negligible effect on the tumor volume compared to other parameters, such as the carrying capacity and proliferation rate. The parameter ranges used in the sensitivity analysis were selected to capture the full range of possible outcomes, from scenarios where the treatment has no effect on the tumor to those where the tumor is eliminated. Even within these wide parameter ranges, *α* and *β* remained less influential on the final tumor volume. However, in the linear treatment model, where the effect of treatment does not diminish over time, *α* becomes the most influential parameter in determining the final tumor volume. This outcome is expected, given that the range of *α* used in both models is the same, and the inclusion of decay in the exponential model reduces the impact of *α* on tumor volume.

Next, we employed the linear model to predict tumor volume at the third time point using three different prediction scenarios: leave-one-out prediction, mouse-specific prediction, and a hybrid group-informed, mouse-specific prediction. Of these methods, the best-performing method was the hybrid method, with an average CCC of 0.85 ± 0.09. This method was able to successfully differentiate between treatment responders and non-responders by propagating the model forward with an average accuracy of 81.26 ± 19.03% across all treatment scenarios. In more detail, the leave-one-out method (Figure 10) correctly predicted only 1 out of 12 responders and 33 out of 37 non-responders, while the mouse-specific method (Figure 11) performed better in predicting responders (12/12) but only correctly identified 20 out of 37 non-responders. The hybrid group-informed, mouse-specific prediction method (Figure 12) was able to identify 11 out of 12 responders and 26 out of 37 non-responders, highlighting that the effect of the treatment indeed differs with subsequent administrations. In the hybrid method, by splitting up the effects of the treatment based on individual time points (i.e., splitting *α* into *α*_1_ and *α*_2_), we isolate and evaluate the effect of combining treatments. To obtain the median resistivity (i.e., 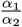) we first calibrate the model using the entire dataset (Figure 8), and then apply this median value when making mouse-specific predictions to determine the individual *α*_2_ for each mouse. Physiologically, this enables the development of a preliminary representation of the effectiveness of an individual dosing event (i.e., a specific administration interval of treatment) within the context of a larger treatment regimen. Combination therapies involving stromal-targeting drugs (i.e., Treatments 2-5) display a higher level of resistivity to treatment than Treatment 1, which uses only NGC (Figure 9). Since our model does not distinguish tumor volume between cancer cells and stroma, we hypothesize that the observed increase in resistivity may (in part) be attributed to stroma depletion caused by these drugs in the first treatment interval rather than a decrease in cancer cell count. Notably, our analysis indicates that the first delivery of treatment is between 0.72 and 4.5 times more effective than the second delivery of treatment. This indicates that there is a diverse level of resistance to treatment, dependent on the specific cocktail of therapies.

Our study contributes to the growing field of mathematical modeling in pancreatic cancer by developing a treatment-agnostic model that integrates multiple chemotherapy and stromal-targeting protocols and performs detailed sensitivity analyses to predict tumor response. This approach builds on the work of Hu et al. [21], who employed a logistic growth assumption in their model of pancreatic cancer dynamics to explore the impact of immunotherapy on tumor progression. While Hu et al. focused on the interactions between tumor cells and the immune system, our model extends these ideas by evaluating the combined effects of various treatment protocols on tumor dynamics, including chemotherapy and stromal-targeting therapies. Additionally, our work complements that of Bratus et al. [20], who explored the evolutionary dynamics of pancreatic cancer cells with a focus on genetic mutations and immune interactions. Whereas their model provides insights into tumor progression through genetic evolution, our model is designed to predict treatment outcomes and guide therapeutic strategies. Furthermore, in response to the gaps highlighted by Dogra et al. [17], who pointed out the scarcity of mathematical models in pancreatic cancer, our study addresses the need for flexible models that can be adapted to different clinical scenarios. Lastly, while Chen et al. [36] developed a PK/PD model focusing on the mechanistic behavior of gemcitabine, our treatment-agnostic approach allows for broader application across various therapeutic combinations, offering a framework for understanding how different treatment strategies can influence pancreatic cancer outcomes.

While our model shows accuracy and practical utility, there are several opportunities for improvement. First, our study modeled all chemotherapy drugs as a single entity, potentially overlooking specific interactions and efficacies of individual drugs. Future work should extend the model to account for the specific treatment protocols of each drug. For instance, in a previous study, we modeled six treatment protocols with two drugs, carefully calibrating each using multiple measurements to capture individual and combined drug effects on breast cancer [37]. However, in this study, we chose a treatment-agnostic approach due to limited data points (three) and sought to assess whether the model could still capture overall tumor development trends without detailed protocol-specific information. Therapies for pancreatic cancer exhibit significant variation, particularly in their mechanisms of action and modes of application [38]. Thus, modifications to the mathematical framework—such as adding or removing parameters or adjusting the prior Bayesian distributions of existing parameters based on the specific mechanisms of each therapy—may improve model fit and enhance predictive power. Another area for improvement is the inclusion of pharmacokinetics in the model to account for drug delivery and distribution, which could provide deeper insights into treatment dynamics (as demonstrated in our previous work in breast cancer [39]). Even though post-processing of the calibration results—by separating the calibration of responders and non-responders and comparing the distributions of the parameters—revealed that non-responders tend to have higher proliferation rates and lower death rates due to treatment (Figure 7), these differences are not accounted for in our current model. Incorporating a data assimilation approach, as demonstrated in [40], could improve our model’s ability to dynamically update predictions based on new data. With the data assimilation approach, our model could be enhanced by differentiating between responders and non-responders earlier in the treatment process, thereby adapting the model parameters in real time to reflect these differences in proliferation and treatment response rates. This would allow for more personalized predictions and potentially more effective treatment strategies, ultimately improving the model’s predictive accuracy. Improvements in the model would, of course, require an increase in the amount and kind of data required to calibrate the resulting model. Increasing the number of mice per treatment group would provide more responders and non-responders, aiding in better differentiation between these populations. It is important to note that in Treatments 4 and 5, every mouse is a responder; however, there are only four mice in Treatment 4 and eight mice in Treatment 5. Increasing the sample size in these groups would offer a more comprehensive understanding of the treatment effects. Furthermore, increasing the number of longitudinal data points beyond the three measurements currently available would improve the model’s predictive power and reliability.

Our mathematical model effectively captures the dynamic progression and regression in a GEM model of pancreatic cancer treated with a range of chemotherapies (cisplatin, paclitaxel, and gemcitabine), both with and without stromal-targeting drugs (calcipotriol and losartan) and an immune checkpoint inhibitor (anti-PD-L1). Additionally, the model successfully predicts tumor volumes through several prediction schemes: mouse-specific predictions (CCC = 0.75 ± 0.05), leave-one-out predictions (CCC = 0.74 ± 0.14), and mouse-specific group-informed predictions (CCC = 0.85 ± 0.09). This work provides a rigorous mathematical framework for characterizing combination therapies for pancreatic cancer, particularly highlighting the interactions between chemotherapies, stromal-targeting drugs, and immunotherapy. By accurately predicting responders and non-responders, the model can help tailor treatment strategies to individual patients, potentially improving therapeutic outcomes. Additionally, the model’s ability to simulate various treatment regimens offers a valuable tool for exploring new combination therapies and optimizing existing ones.

## Acknowledgements

We thank the American Cancer Society Grant RSG-18-006-01-CCE and the National Institutes of Health for funding via R01CA240589, R01CA276540, 1U24CA231858, and U24CA226110. We thank the Cancer Prevention and Research Institute of Texas for support through CPRIT RR160005. T.E.Y. is a CPRIT Scholar in Cancer Research. We also thank the Texas Advanced Computing Center for providing high-performance computing resources.

## Author Contributions

EABFL, TEY conceptualized and supervised the study. KV, EABFL developed the computational method, framework, and analysis. RZ developed the experimental data and physiological motivation for models. HC, MG conducted the enrollment, treatment, and tumor size measurements. KV, EABFL designed the ODEs and model framework. KV performed Bayesian model calibrations, sensitivity analysis, and other computations. KV, EABFL wrote the initial draft of the manuscript. All authors revised and approved the manuscript.

## Data availability

The datasets generated and/or analyzed during the current study are available upon request.

## Code availability

Our code is shared publicly on GitHub upon publication of this work and can be found at https://github.com/krithikvishwanath/NGC_Therapy_Modeling.

## Supplementary Material

**Supplemental Figure 1.**
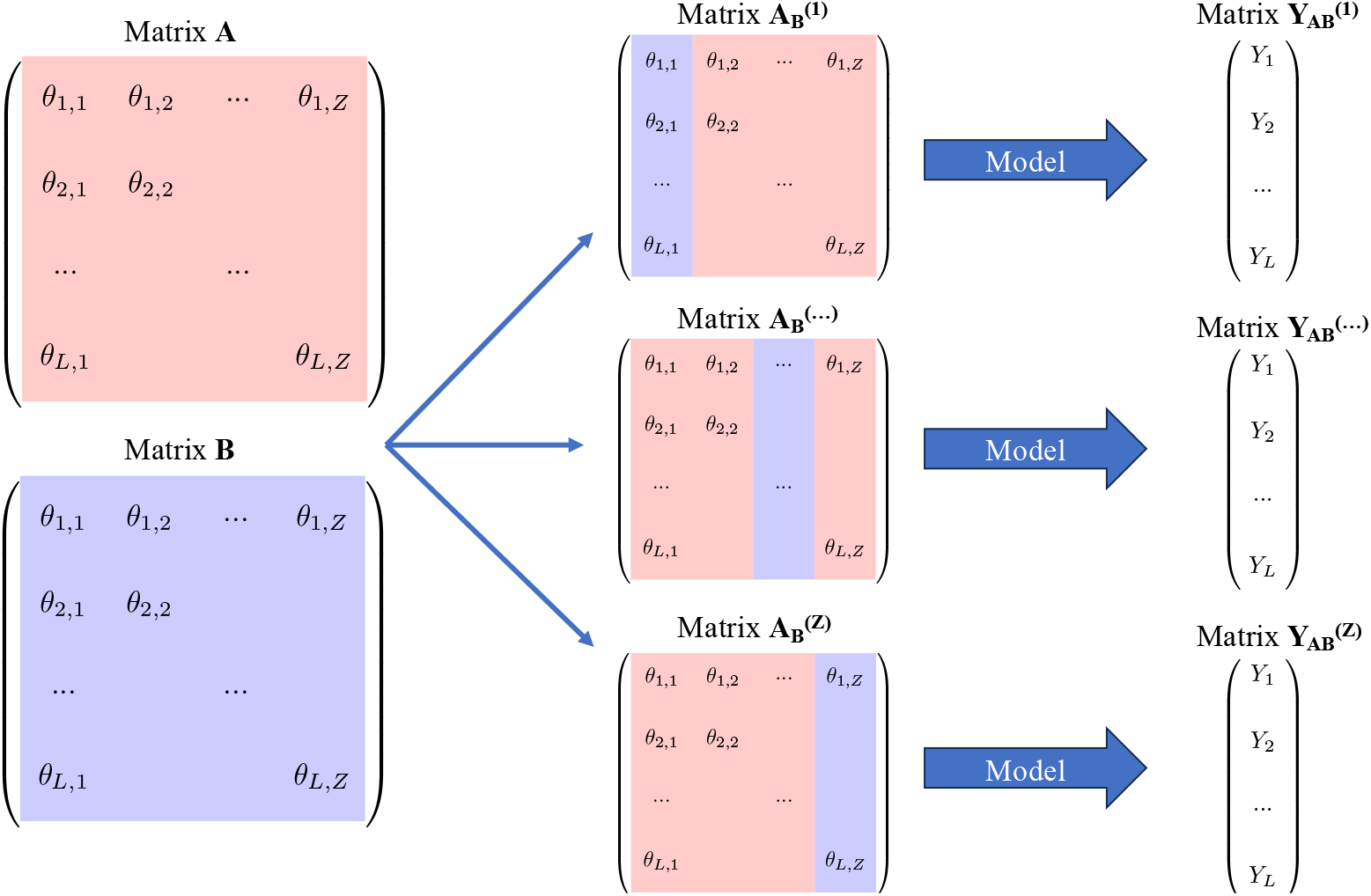
Visual representation of total sensitivity index estimation using matrix notion. Theta, ***θ***, is defined as a vector of parameters for a particular model. We began by developing matrices ***A*** and ***B***, where arbitrary column *z* is a vector of *L* values randomly generated from the sample space of the *z*^*th*^ parameter. ***A*** and ***B*** are then used as components of a new hybrid matrix, with one column from ***B*** and the rest from ***A***. Then, the model is propagated for each hybrid matrix and for matrix ***A***. Variations between the output of matrix ***A*** compared to each hybrid matrix allow insight of parameter influence on model output, within the sampling space.

